# Microbiome and climate: Skin microbial diversity and community functions of *Polypedates megacephalus* (Anura: Rhacophoridae) associated with bioclimate

**DOI:** 10.1101/2024.09.16.613297

**Authors:** Dan Sun, Yewei Liu, Shipeng Zhou, Madhava Meegaskumbura

## Abstract

The microbiome inhabiting animal skin plays a crucial role in host fitness by influencing both the composition and function of microbial communities. Environmental factors, including climate, significantly impact microbial diversity and the functional attributes of these communities. However, it remains unclear how specific climatic factors affect amphibian skin microbial composition, community function, and the relationship between these two aspects. Given that amphibians are poikilotherms, and thus more susceptible to temperature fluctuations, understanding these effects is particularly important. Here, we investigated the skin microbiome of the rhacophorid tree frog *Polypedates megacephalus* across different climatic regimes using 16S rRNA gene sequencing. Skin swab samples were collected from nine populations of *P. megacephalus* adults in the Guangxi region, China. The majority of the core microbiota were found to belong to the genus *Pseudomonas*. Our findings indicate that microbial community diversity, composition, and function are associated with changes in climatic conditions. Specifically, the taxonomic and functional diversity of the skin microbiome increased in response to greater climate variability, particularly in temperature fluctuations. Additionally, the functional attributes of microbial communities changed in parallel with shifts in community diversity and composition, suggesting that environmental filtering driven by climate changes negatively impacts microbial community functional redundancy. These results highlight the critical influence of climatic factors on amphibian skin microbiomes and offer new insights into how microbial composition and function contribute to host adaptation in varying environmental conditions.

**IMPORTANCE:** This study is important in understanding the association between climate variability, microbial diversity, and host adaptation in amphibians, particularly vulnerable to environmental changes due to their poikilothermic nature. Amphibians rely on their skin microbiome for key functions like disease resistance, yet little is known about how climate fluctuations impact these microbial communities. By analyzing the microbiome of *Polypedates megacephalus* across different climatic regimes, our analysis reveals that while climate variability enhances microbial diversity, it reduces functional redundancy. These findings highlight the potential ecological consequences of climate change and emphasize the need to integrate microbiome health into amphibian conservation strategies.

## INTRODUCTION

The microbiome inhabiting host skin plays a key role in host fitness by altering their community composition and function (1-3). Many studies have shown that changes in microbiome diversity or dominant bacterial communities have a negative impact on the health of human, wild, and domesticated animals (4-8). The characteristics of skin microbiota can be shaped by biotic factors (e.g., host-related traits) and abiotic factors (e.g., temperature and water quality) (9). In particular, climate-related parameters have been highlighted as one vital element for understanding ecology and evolution of host-associated microbiome (10).

Amphibians serve as the crucial components in both aquatic and terrestrial ecosystems, with their significance extending even to human health (11, 12). The exposed skin of amphibians, as the key component determining respiration, thermoregulation, osmoregulation, pigmentation, and protection from predators and pathogens, exhibits extremely sensitive to environmental variations (13). The skin microbial community compositions and interactions play an important role in the dynamics of infectious diseases (14, 15), such as chytridiomycosis defined as the most destructive amphibian skin disease (16).

Several studies show the relationships between bioclimate and amphibian skin microbial community (10, 17-20); for example, skin microbiota exhibited more diverse in the colder and less stable temperature conditions compared with those in warm and less temperature variations (10). Nevertheless, Bacigalupe et al. (21) did not find the significant relationship between skin bacterial communities of the four-eyed frog (*Pleurodema thaul*) and climate factors. The different interpretations for the effects of ecological factors on the structure and compositions of skin bacterial diversity are dependent on different scale-based measurements (e.g., biogeographic gradients and host species) (22). Given the influence of host-specific and biogeographic area on host skin microbiome, it is necessary to in-deep study the structure and compositions of skin bacterial community under the context of various climatic conditions. This will not only improve the understanding how climate factors influence skin microbiota, but can predict the adaptation of amphibians to the rapid changes in climate.

Environmental conditions, including climate, not only affect microbial diversity but have profound effects on the functional attributes of the microbial community (23, 24), while few studies have considered the functional attributes of amphibian skin microbial community under the context of climatic regimes (19). On the other hand, environmental changes are able to affect the relationships between microbial community composition and functional attributes due to the different degrees of influences of environmental filtering on microbial functional redundancy (25), where multiple distinct taxa have similar functions (26). However, much less known is about the effects of climatic variations on the relationships between amphibian skin microbial community and their functions.

*Polypedates megacephalus*, an Old-World tree frog (family Rhacophoridae), is widespread in southern China and Southeast Asia, occupying a broad niche space (27). Given the species’ wide geographic range and the significant impact of climate on animal microbiome, we hypothesized that the skin microbiota of *P. megacephalus* is closely linked to climatic factors. To test this, we sampled skin microbiota from populations inhabiting different sites along a latitudinal gradient in Guangxi, China, characterized by varying environmental conditions. We predicted that microbial community diversity and composition would be strongly influenced by climate-related factors and that the diversity and composition of microbial community functional attributes would show correlations (positive or negative) with climatic characteristics. Additionally, we anticipated that microbial community functional attributes and taxonomic diversity would exhibit similar responses to climatic variations, reflecting the impact of climate fluctuations on microbial functional redundancy.

## RESULTS

The taxonomic composition of OTUs on *P. megacephalus* frogs skin bacteria predominately consisted of the phyla Proteobacteria, Bacteroidota, Actinobacteriota, Cyanobacteria, Planctomycetota, and Firmicutes, with the relative abundance of 69.76%, 6.89%, 5.33%, 3.68%, 2.84%, and 2.47%, respectively. Hierarchical clustering of microbiome revealed that *P. megacephalus* skin had a site-specific microbial community structure (Fig. 1).

**Figure 1.**
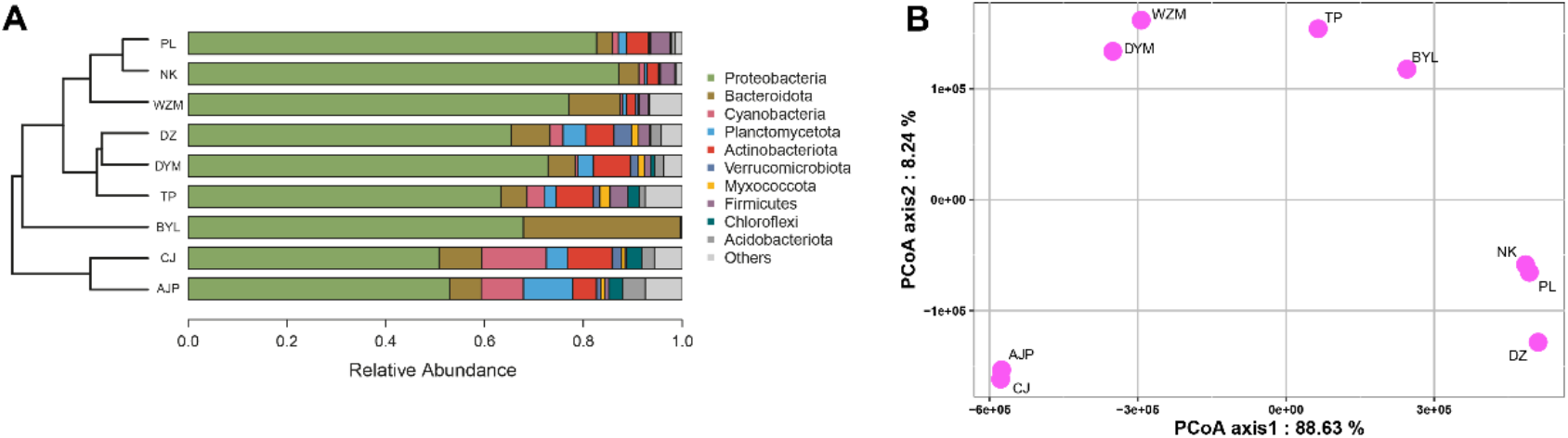
UPGMA and stacked bar of skin bacteria at the phylum level. The composition of bacteria showed different for *P. megacephalus* populations across sampling sites (A). The PCoA scatterplots show the distribution of sampling sites based on Euclidean distance (B). The abbreviations of these geographic sites: Anjiangping (AJP), Cujiang (CJ), Biyun Lake (BYL), Tianping (TP), Dayao Mountain (DYM), Dongzhong (DZ), Wuzhi Mountain (WZM), Nakuan (NK), Pinglong (PL).

Total 56 OTUs were was identified as a core bacterial community that dominated skin bacteriome of most frogs (Table 1). The core community exhibited 46% average relative abundance (minimum 8%, maximum 96%). The majority of core OTUs were identified as belonging to the genus Pseudomonas, which accounted for 10.20% of average relative abundance per frog. One individual with the lowest core community abundance (3%) was dominated by the members of the family Pasteurellaceae.

**Table 1.** Taxonomy of core microbiome present on 90% of *P. megacephalus* frogs sampled.

| OUT_ID | Taxonomy | Average relative abundance (%) |
| --- | --- | --- |
| 1, 408, 1207, 7097, 8613, 13995 | p__Proteobacteria, c__Gammaproteobacteria, o__Pseudomonadales, f__Pseudomonadaceae, g__Pseudomonas | 12.24 |
| 2, 94, 1075, 1174, 1201, 9030, 9732, 11586, 11850, 12242, 12717, 12839 | p__Proteobacteria, c__Gammaproteobacteria, o__Gammaproteobacteria, f__Comamonadaceae | 10.36 |
| 3 | p__Proteobacteria, c__Gammaproteobacteria, o__Enterobacterales, f__Enterobacteriaceae, g__Cedecea | 9.19 |
| 4 | p__Proteobacteria, c__Gammaproteobacteria, o__Burkholderiales, f__Comamonadaceae, g__Comamonas | 1.67 |
| 8 | p__Proteobacteria, c__Gammaproteobacteria, o__Enterobacterales, f__Hafniaceae, g__Hafnia-Obesumbacterium | 1.18 |
| 11 | p__Proteobacteria, c__Gammaproteobacteria, o__Burkholderiales, f__Oxalobacteraceae, g__Janthinobacterium | 0.90 |
| 14 | p__Proteobacteria, c__Gammaproteobacteria, o__Xanthomonadales, f__Xanthomonadaceae, g__Stenotrophomonas | 0.61 |
| 15 | p__Firmicutes, c__Bacilli, o__Exiguobacterales, f__Exiguobacteraceae, Exiguobacterium | 0.91 |
| 16, 36, 37 | p__Proteobacteria; c__Alphaproteobacteria; o__Rhizobiales; f__Rhizobiaceae; g__Allorhizobium-Neorhizobium-Pararhizobium-Rhizobium | 1.50 |
| 17 | p__Proteobacteria, c__Gammaproteobacteria, o__Burkholderiales, f__Comamonadaceae, g__Pseudorhodoferax | 1.30 |
| 19 | p__Actinobacteriota, c__Actinobacteria, o__Micrococcales, f__Microbacteriaceae, g__Microbacterium | 0.76 |
| 23, 121 | p__Proteobacteria, c__Alphaproteobacteria, o__Rhizobiales, f__Devosiaceae, g__Devosia | 0.49 |
| 26 | p__Proteobacteria, c__Gammaproteobacteria, o__Burkholderiales, f__Oxalobacteraceae, g__Herbaspirillum | 0.44 |
| 28 | p__Proteobacteria, c__Alphaproteobacteria, o__Rhizobiales, f__Beijerinckiaceae, g__Bosea | 0.30 |
| 31 | p__Proteobacteria, c__Gammaproteobacteria, o__Pseudomonadales, f__Moraxellaceae, g__Enhydrobacter | 0.22 |
| 48 | p__Proteobacteria, c__Alphaproteobacteria, o__Rhizobiales, f__Xanthomonadaceae, g__Bradyrhizobium | 0.39 |
| 52 | p__Proteobacteria, c__Gammaproteobacteria, o__Pseudomonadales, f__Moraxellaceae, g__Acinetobacter | 0.28 |
| 60 | Proteobacteria, c__ Gammaproteobacteria, o__ Burkholderiales, f__ Oxalobacteraceae, g__ Duganella | 0.13 |
| 73 | p__ Proteobacteria, c__ Gammaproteobacteria, o__ Burkholderiales, f__ Burkholderiaceae, Burkholderia-Caballeronia-Paraburkholderia | 0.22 |
| 74 | p__ Actinobacteriota, c__ Actinobacteria, o__ Micrococcales, f__ Micrococcaceae, g__ Arthrobacter | 0.12 |
| 83 | p__ Proteobacteria, c__ Alphaproteobacteria, o__ Rhizobiales, f__ Beijerinckiaceae | 0.18 |
| 92 | p__ Proteobacteria, c__ Alphaproteobacteria, o__ Sphingomonadales, f__ Sphingomonadaceae, g__ Hephaestia | 0.14 |
| 100 | p__ Proteobacteria, c__ Gammaproteobacteria, o__ Enterobacterales, f__ Erwiniaceae | 0.14 |
| 102 | p__ Actinobacteriota, c__ Actinobacteria, o__ Micrococcales, f__ Intrasporangiaceae | 0.10 |
| 147 | p__ Proteobacteria, c__ Alphaproteobacteria, o__ Rhizobiales, f__ Rhizobiaceae, g__ Pseudochrobactrum | 0.09 |
| 156 | p__ Actinobacteriota, c__ Actinobacteria, o__ Micrococcales, f__ Microbacteriaceae, g__ Leifsonia | 0.09 |
| 402 | p__ Proteobacteria, c__ Alphaproteobacteria, o__ Sphingomonadales, f__ Sphingomonadaceae, g__ Novosphingobium | 0.05 |
| 571, 808, 10525 | p__ Proteobacteria, c__ Alphaproteobacteria, o__ Sphingomonadales, f__ Sphingomonadaceae, g__ Sphingomonas | 0.28 |
| 663 | p__ Actinobacteriota, c__ Actinobacteria, o__ Micrococcales, f__ Microbacteriaceae, g__ Galbitalea | 0.35 |
| 727 | p__ Proteobacteria, c__ Gammaproteobacteria, o__ Burkholderiales, f__ Oxalobacteraceae, g__ Massilia | 0.52 |
| 1628 | p__ Actinobacteriota, c__ Actinobacteria, o__ Micrococcales, f__ Cellulomonadaceae, g__ Cellulomonas | 0.04 |
| 5256 | p__ Proteobacteria, c__ Gammaproteobacteria, o__ Enterobacterales, f__ Enterobacteriaceae | 0.13 |
| 10178 | p__ Proteobacteria, c__ Gammaproteobacteria, o__ Burkholderiales, f__ Comamonadaceae, g__ Acidovorax | 0.28 |
| 13900 | p__ Proteobacteria, c__ Gammaproteobacteria, o__ Enterobacterales | 0.51 |
| 13990 | p__ Proteobacteria, c__ Gammaproteobacteria, o__ Enterobacterales, f__ Yersiniaceae | 0.16 |

Skin microbial alpha diversity indices had significant relationships with climatic factors (Mantel test: Pearson’s *r* = 0.162, *P* = 0.024; Fig. 2). The frogs exhibited increased skin microbial richness and phylogenetic diversity in colder climate or drier climate (Fig. S1). For instance, skin microbiota showed a decrease of alpha diversity with the minimum temperature of coldest month (Bio6) (richness: *R*^*2*^ = 0.526, *p* < 0.0001; Shannon: *R*^*2*^ = 0.469, *p* < 0.0001; PD: *R*^*2*^ = 0.319, *p* = 0.001).

**Figure 2.**
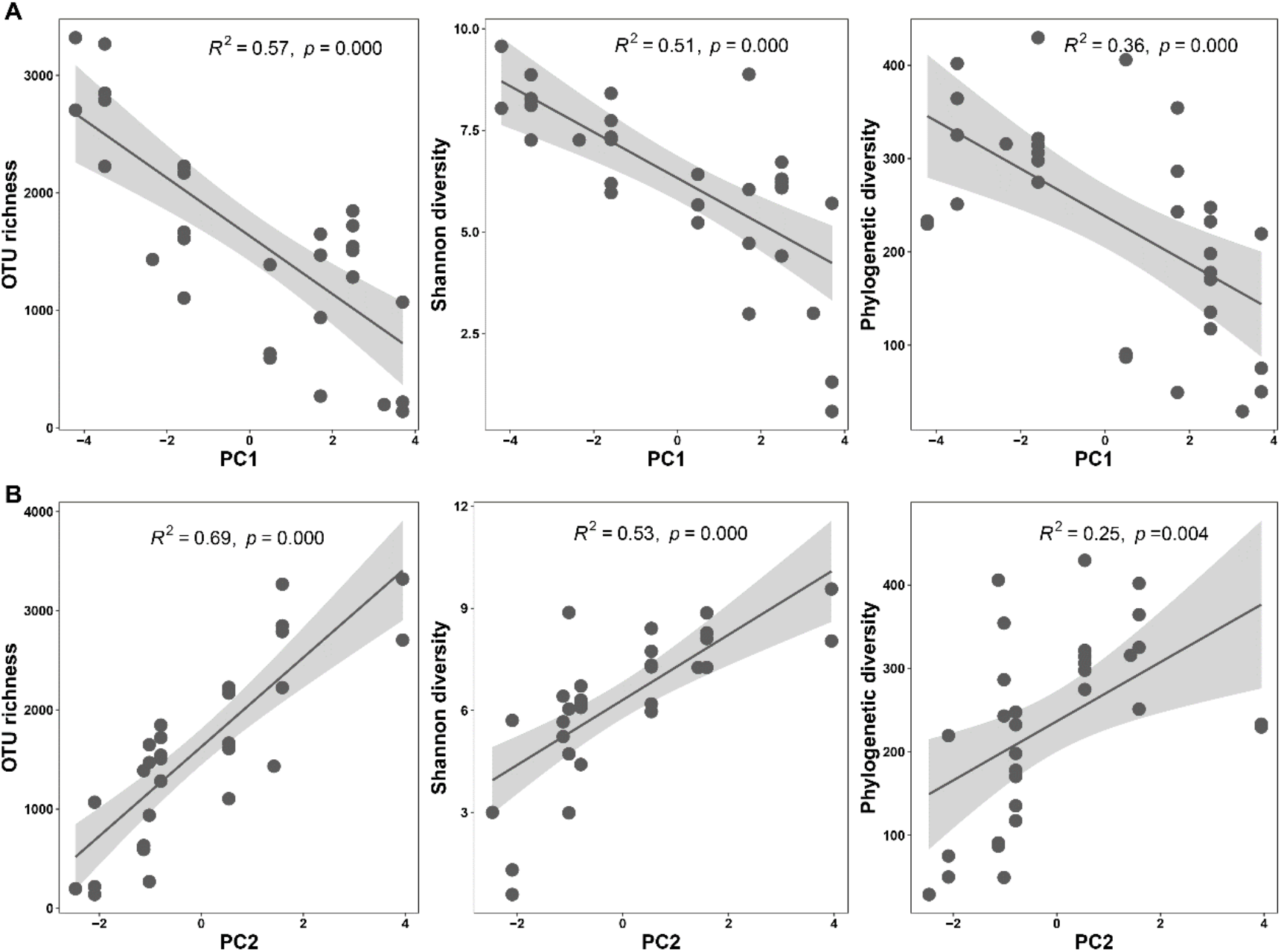
Linear regression plot between microbial alpha diversity with temperature-related factor (A) and precipitation-related factor (B). Gray shading represents 95% Confidence Interval.

The microbial beta-diversity had significant associations with both temperature factor (PC1; *F* = 3.17, *R*^*2*^ = 0.10, *p* < 0.0001) and precipitation factor (PC2; *F* = 2.01, *R*^*2*^ = 0.06, *p* = 0.006) (Fig. 2A and 2B), consistent with the result of Mantel test (Pearson’s *r* = 0.234, *p* = 0.001).

We found that the members of the Beijerinckiaceae, Xanthobacteraceae, Burkholderiaceae, Deinococcaceae, Oxalobacteraceae, Sphingomonadaceae, and unidentified_Chloroplast families correlate with temperature factor, which increased with colder climate; the members of Comamonadaceae, Moraxellaceae, Bdellovibrionaceae, Exiguobacteraceae, Hafniaceae, Microbacteriaceae, Sphingobacteriaceae, and Xanthomonadaceae families were correlated with precipitation factor (Fig. 3C).

**Figure 3.**
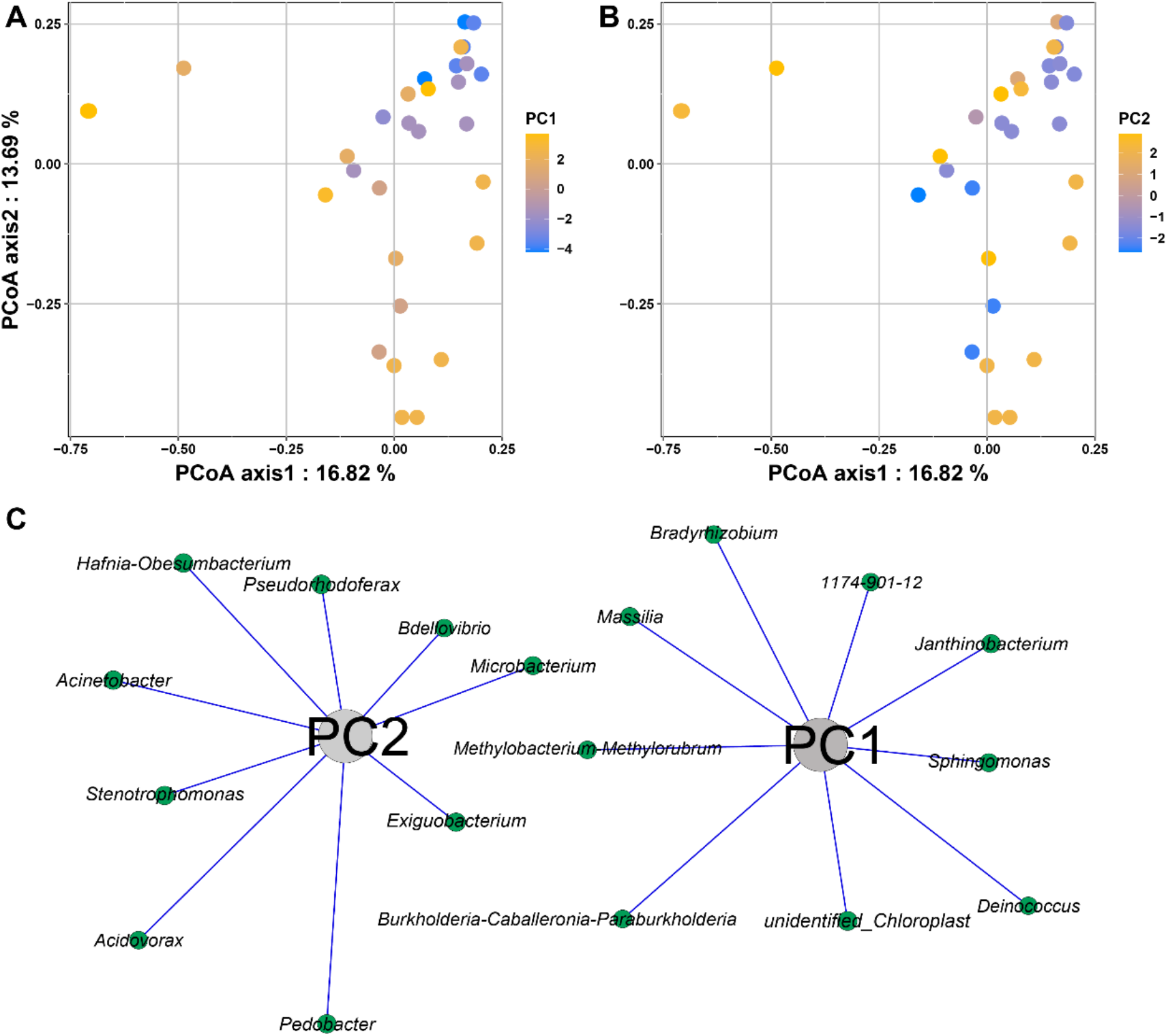
PCoA scatterplots show the variations in microbial community composition based on Bray-Curis distance (A and B) and the correlations of bacterial genus with climatic indices (Spearman’ *r* > |0.75|, *p* < 0.01) (C).

Most functional features were associated to metabolism (48.65% ± 0.019%). The top five functional categories in the KEGG pathways (level 2 KOs) included membrane transport (12.81% ± 0.018%), amino acid metabolism (10.25% ± 0.006%), carbohydrate metabolism (9.75% ± 0.003%), replicate and repair (6.56% ± 0.003%), energy metabolism (5.46% ± 0.005%).

The climatic factors significantly influenced the functional diversity of skin microbiome of the tree frogs (Fig. 4), with the change tendency similar to microbial diversity under the context of PC1 and PC2 (Fig. 1).

**Figure 4.**
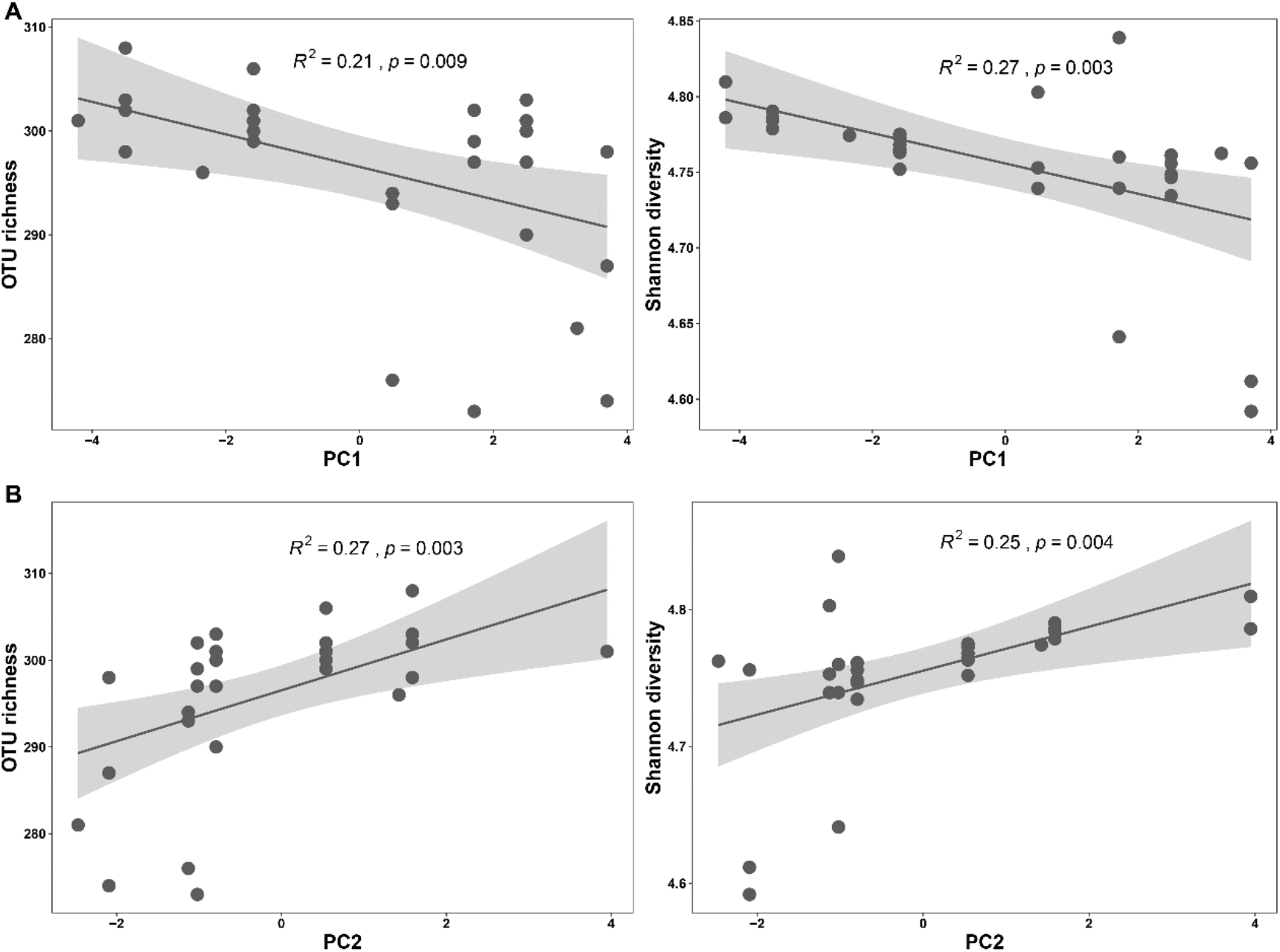
Linear regression plot between microbial functional diversity and temperature-related factor (A) and precipitation-related factor (B). Shading represents 95% Confidence Interval.

The variations in skin microbial function compositions significantly correlated with bioclimatic variables in Mantel test (Spearman’ *r* = 0.150, *p* = 0.031); this pattern also found in the evaluation of the associations of functional dissimilarity with temperature-related factor (*F* = 4.789, *R*^*2*^ = 0.142, *p* < 0.006) and precipitation-related factor (*F* = 4.029, *R*^*2*^ = 0.122, *p* = 0.021) in the permutational multivariate analysis of variance (PERMANOVA).

Many of functional items positively related to climate-related factors. For example, the relative abundance of ATP-binding cassette (ABC) transporters had an increased trend with the minimum temperature of coldest month (Bio6); ABC transporters had a decreased variation with mean monthly precipitation amount of the coldest quarter (Bio19) (Fig. 5).

**Figure 5.**
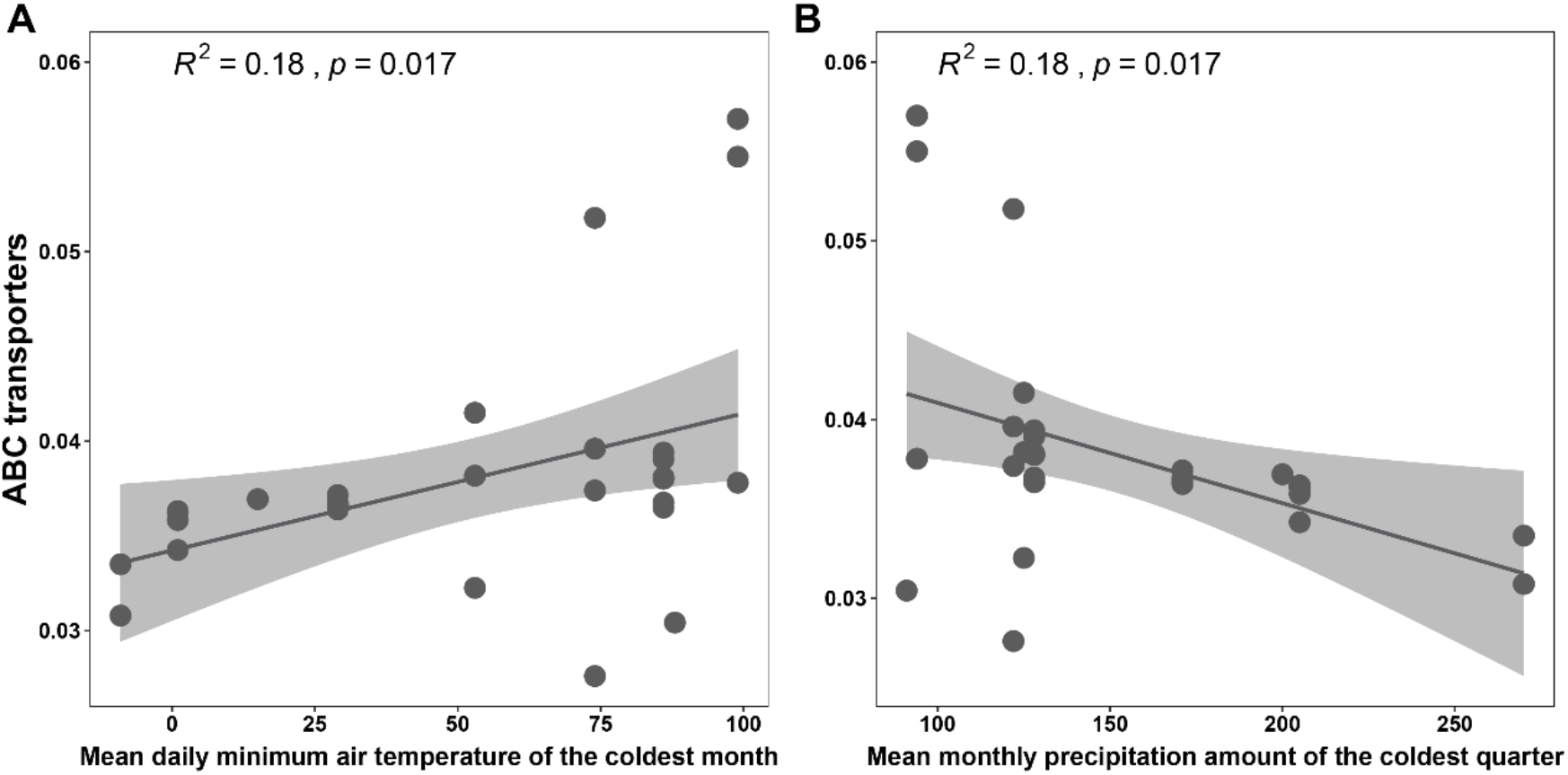
Linear regression between ABC transporters and the minimum temperature of the coldest month (bio6) and mean monthly precipitation amount of the coldest quarter (bio19).

We found that skin microbial community diversity positively correlated with the community functional diversity (Shannon: *R*^*2*^ = 0.75, *P* < 0.001) (Fig. 6A). This pattern was also observed between microbial community compositions and functional attributes (linear regression: *R*^*2*^ = 0.41, *P* < 0.001; Mantel test: *r* = 0.7437, *P* < 0.001) (Fig. 6B).

**Figure 6.**
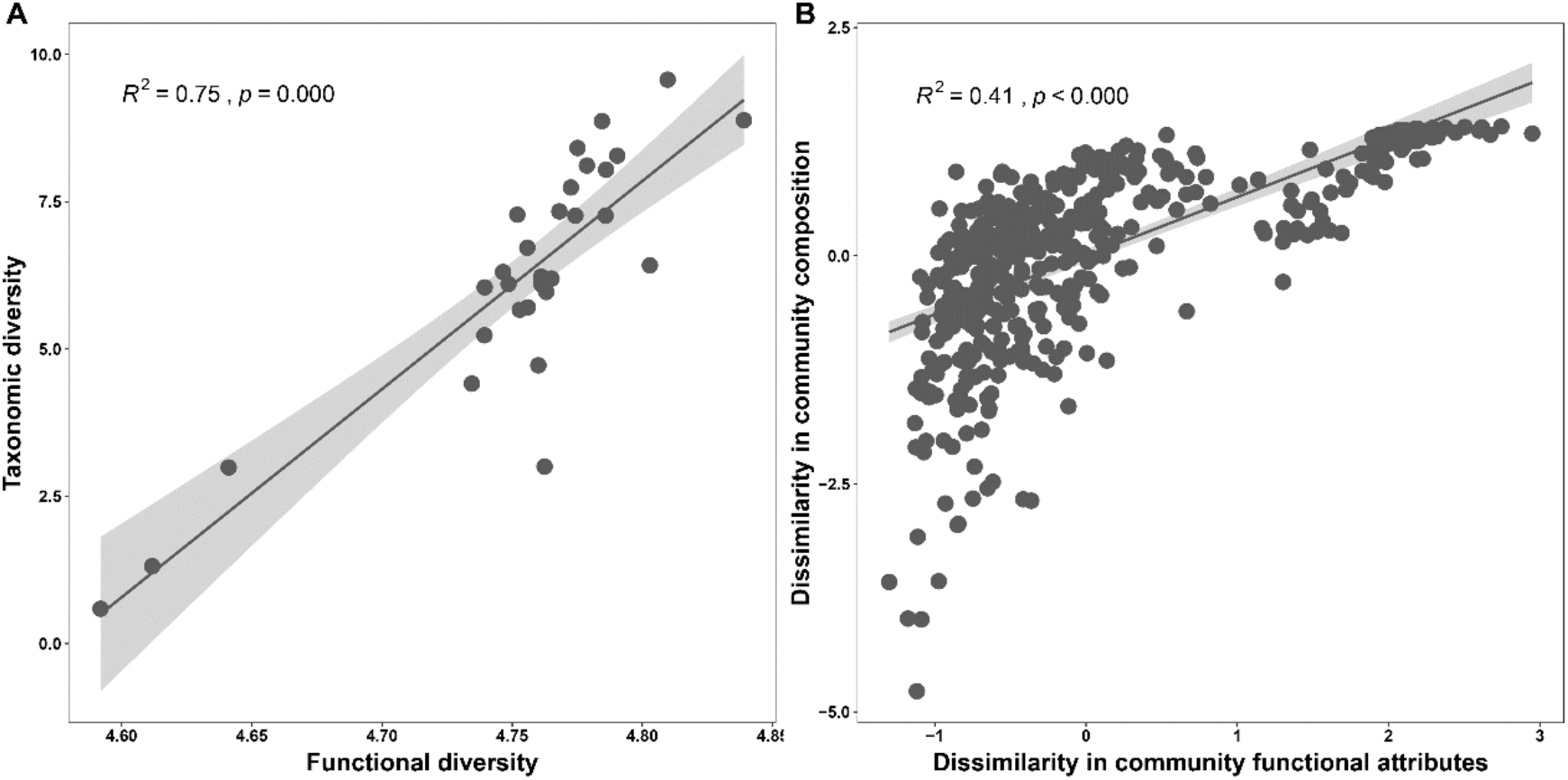
A linear relationship between community functional Shannon diversity and taxonomic Shannon diversity (A). A linear relationship between dissimilarities of microbial community composition and functional attributes (B).

## DISCUSSION

We found that the skin microbial diversity and composition as well as community function of *P. megacephalus* frogs associate with changes of climatic conditions. The taxonomical diversity and functional diversity of the skin microbiome increased with fluctuations in climate, particularly regarding temperature variations. Furthermore, the functional attributes of the microbial community synchronously change community diversity and composition. This suggests that environmental filtering related to climate changes exerts negative influences on microbial community functional redundancy, with a cold-wet climate could result in a lower value in microbial community functional redundancy.

The phylum Proteobacteria constituted the highest abundance in the skin microbiota harbor *P. megacephalus* frogs, as previously observed in temperate and tropical amphibian populations (28-30), and that the relative abundance of this phylum positively increased with colder climate (10). The rich Proteobacteria might aid in the conversion of chitin to chitosan and cellulose (31). Despite *P. megacephalus* skin showing a site-specific microbial community structure, microbiota from certain sites were not clustered together with those from proximal sites. For instance, the microbiota from DZ location was more similar to DYM rather than those (PL and NK locations) relatively close distance to DZ. The result suggests that geographic distances did not have a strong effect on the differences in microbiota due to the effects from local habitats (32).

We found that the alpha diversity of skin microbiome related to climatic factors, with increased diversity along with the changing trend towards the cold and wet climate. The lower diversity of skin microbiome in warm climates might be attributed to certain microbes that have higher thermal growth optima, making them intensified competitive interactions in stable thermal environments. For instance, the genus *Acidovorax*, with their relative abundance decreased in cold climate; this might lead to the abundance of other taxa have similar niche increased owning to the requirements of nutrients or growth conditions (10, 33). Another important point is that certain microbes have stronger dependence on wet climate. For instance, we found the members of the families of Comamonadaceae and Moraxellaceae highly related to precipitation factor; these families were enriched in Panama amphibians (2). Conversely, a greater microbial diversity of *P. megacephalus* in cold climate may indicate that hosts have evolved more complex adaptive immunity (18, 20).

For skin microbial community beta diversity, our results showed that temperature-related parameters could drive the dissimilarities of the skin microbial community more than that of precipitation-related parameters. One possible explanation for this result is that temperature may strongly affect the colonization of certain microbes, leading to changes in the structure and compositions of a microbial community (34). Despite site climate difference, we identified some core taxa of the white-spotted thigh treefrog *P. megacephalus*, particularly members of the genus *Pseudomonas*, may possess vital functions for hosts, such as pathogens resistance (35, 36). These results suggest that certain skin microbiota tend to show an effective response in facing of various climate conditions, to facilitate hosts harbour in a broader geographic area.

The previous study showed that microbial metabolic processes are strongly affected by natural fluctuations in temperature and precipitation (37). The identity of the metabolic features under the effects of climate shed some light on the possible mechanisms for changes in the structure and compositions of microbial communities and hosts coping with climate variations. For instance, the relative abundance of ABC transporters increased with the temperature of the coldest month and decreased with precipitation of the coldest quarter; these relationships are similar to the finding that the ABC transporters of coqui frogs (*Eleutherodactylus coqui*) microbiome metabolic activities were enriched during the cool-dry season (37); because the cooler climate suppress host immunity, leading to the increased critical functions of ABC transporters including nutrient import and molecule export (38, 39).

The community functional traits are positively related to cold climate, showing a broader distribution; this relationship is a similar tendency to the variations in microbial taxonomical diversity under climate fluctuations. Furthermore, microbial community functional attributes synchronously changed community compositions. The results indicate that environment filtering with regard to the variations in climatic conditions weakens microbial community functional redundancy due to the cold climate may drive amphibian hosts to select cold-tolerant microbe, reducing community functional redundancy (25, 40). Skin microbial community composition and the distribution of functional traits are essential for amphibians to cope with the changes in climate conditions. Accordingly, further study is needed to enhance the understanding of the relationships of community functions with taxonomical diversity and climate changes by extending sampling sites for *P. megacephalus* and amphibian species.

Climate change such as higher temperature and heavier rainfall events would dramatically affect microbial diversity and function (41, 42). The temperature-dependent and precipitation-dependent patterns of microbial community composition and function in *P. megacephalus*, suggest that climate change will decrease the diversity of host skin microbiome. Importantly, the altered microbiome structure may impact host immunity and performance, even the persistence of local *P. megacephalus* populations. In addition, our study provides new insights into the compositions of core bacteria and climate-related taxa of *P. megacephalus*, contributing to the amphibian conservation through microbiota-associated strategies. However, broader geographic sampling and taxonomical sampling of natural species and populations is needed to advance the understanding of climate effects on amphibian skin microbial ecology and evolution.

In conclusion, climate influences the skin microbial community composition and functional traits in amphibian populations at different sites. Our results not only advance the understanding of the associations of climatic factors with amphibian skin microbial community diversity and community functional attributes. But contribute to predicting the adaptation of amphibians to the rapid changes in climate. Our results reveal that environmental filtering, which relates to climate, has influences on the diversity and composition of amphibian skin microbial and community functional attributes, as well as their relationships.

## MATERIALS AND METHODS

### Field sample collection

We collected skin swab samples from *P. megacephalus* adult individuals from nine wild sites in 2021 in Guangxi region, China. These sites included Anjiangping (*N*=2), Cujiang (*N*=4), Biyun Lake (*N*=1), Tianping (*N*=6), Dayao Mountain (*N*=1), Dongzhong (*N*=4), Wuzhi Mountain (*N*=3), Nakuan (*N*=3), Pinglong (*N*=7). Epidermal swabs were obtained from all individuals using sterile swabs and subsequently released at their captured locations (43). We followed standardized protocols and biosecurity measures while collecting and analyzing the swabs to prevent cross contamination between individuals and transfer of pathogens across habitats. Ethical clearance was obtained from the Animal Care & Welfare Committee of Guangxi University (GXU2020-501).

### Bioclimatic variables

We downloaded 19 bioclimatic variables from the CHELSA version 1.2 database at a resolution of 30 arcsec (44). These bioclimatic variables consisted of mean annual air temperature (bio1), mean diurnal air temperature range (bio2), isothermality (bio3), temperature seasonality (bio4), mean daily maximum air temperature of the warmest month (bio5), mean daily minimum air temperature of the coldest month (bio6), annual range of air temperature (bio7), mean daily mean air temperatures of the wettest quarter (bio8), mean daily air temperatures of the driest quarter (bio9), mean daily mean air temperatures of the warmest quarter (bio10), mean daily mean air temperatures of the coldest quarter (bio11), annual precipitation (bio12), precipitation amount of the wettest month (bio13), precipitation amount of the driest month (bio14), precipitation seasonality (bio15), mean monthly precipitation amount of the wettest quarter (bio16), mean monthly precipitation amount of the driest quarter (bio17), mean monthly precipitation amount of the warmest quarter (bio18), mean monthly precipitation amount of the coldest quarter (bio19). To reduce the dimensions of climatic factors, we performed principal component analysis (PCA) to consolidate the temperature and precipitation variables, respectively. The first principal component (PC) scores were used as the temperature-related and precipitation-related factors.

### DNA extraction and 16S rRNA gene sequencing

We extracted DNA from the swabs using Qiagen DNeasy Blood and Tissue Kit, which subsequently were amplified for genomic DNA using PCR. The V4 region of 16S rRNA gene was amplified using the specific primers (515F: GTGCCAGCMGCCGCGGTAA and 806R: GGACTACHVGGGTWTCTAAT) with the barcodes. Total 15 µL of Phusion® High-Fidelity PCR Master Mix (New England Biolabs), 0.2 µM of forward and reverse primers, and 10 ng DNA extract were used in each PCR amplification. The conditions for PCR amplification consisted of initial denaturation at 98 °C for 1 min, 30 cycles of denaturation at 98 °C for 10 s, annealing at 50 °C for 30 s, and elongation at 72 °C for 30 s and 72°C for 5 min. The PCR products were purified using magnetic bead purification. Sequencing libraries were generated and indexes were added. The library quality was evaluated using the Qubit® 2.0 Fluorometer (Life Technologies, California, USA) and Agilent 2100 Bioanalyzer system (Agilent Technologies, California, USA). Quantified libraries pooled and were sequenced on an Illumina platform and 250 bp paired-end reads were generated.

### Bioinformatic analysis

The paired-end reads were processed using Python v3.6.13 and adaptors were removed through cutadapt v3.3. The paired-end reads were merged using FLASH v1.2.11 (45). Quality filtering on the raw tags were performed using the fastp v0.23.1 (46). The chimera sequences were removed with the vsearch package (47).

Sequences analysis was performed through Uparse v7.0 (48), and sequences with ≥ 97% similarity were assigned to the same microbial operational taxonomic unit (OTU). Each OTU used the representative sequence was annotated taxonomic information using the silva database based on Mothur algorithm. OTUs abundance were rarefied to 13,573 sequences per sample based on the sample with the least sequences. The multiple sequences were aligned using the MUSCLE v3.8.31 (49), to analyze the phylogenetic relationship of different OTUs. Core skin microbiota was defined as OTUs that were present on at least 90% of all frog samples.

The phylogenetic investigation of communities by reconstruction of unobserved states (PICRUSt) approach can both predict the Kyoto Encyclopedia of Genes and Genomes (KEGG) Ortholog (KO) functional profiles of microbial communities via 16S rRNA gene sequences (50). We used PICRUSt to predict the functional characteristics of microbiota which reside on *P. megacephalus* skin (51).

### Statistical analysis

The general community structure and cluster patterns of microbiome across sites were measured using the unweighted pair-group with arithmetic mean (UPGMA) method based on Bray–Curtis distances. Furthermore, to test whether climatic related factors have strong influences on the taxonomical and community functional diversity and composition of *P. megacephalus* microbiome, we firstly calculated three alpha diversity metrics indices: Shannon’s diversity index, OTU richness, and Faith’s Phylogenetic Diversity (Faith’s PD). Secondly, we calculated the weighted Bray-Curtis and unweighted Jacarrd distances for the microbial beta diversity and function using the adonis2 function in the “vegan” package with 9999 random permutations. Principal coordinate analyses (PCoA) were used to visualize the distance correlations among skin microbial community and function dissimilarity of frog samples. Thirdly, we analyzed the relationships between skin microbial alpha-diversity metrics and beta-diversity as well as community functional attributes (KO) between frog samples and bioclimatic variables using mantel test and linear regression. We analyzed the changes in the microbial community composition in relation to the climatic factors at the genus level using spearman correlation in the “vegan” package (52), to evaluate the significant associations of climatic factors with microbial taxa. To test whether skin microbial community compositions and community functional attributes have similar responses to climatic changes, we analyzed the relationships between microbial community diversity and community functional attributes.

## DATA AVAILABILITY

The 16S sequence data have been uploaded to NCBI with the accession number PRJNA1155111.

## AUTHOR CONTRIBUTIONS

D.S. and M.M. conceived the project. D.S., Y.W.L., and S.P.Z. investigated and collected the skin swab samples. D.S., Y.W.L., and S.P.Z. performed the experiments. D.S., Y.W.L., S.P.Z., and M.M. processed and analyzed the data. D.S. and M.M. wrote the manuscript. All authors consented to the final version of the manuscript.

We declare no competing interests.

## ETHICS STATEMENT

No amphibians were sacrificed, and skin swabbing was employed following safe protocols that do not harm the frogs—ethical approval from the Animal Care & Welfare Committee of Guangxi University (GXU2020-501).

## SUPPLEMENTAL MATERIAL

FIGURE S1. Linear regression plot between microbial alpha diversity with and the minimum temperature of the coldest month (A) and mean monthly precipitation amount of the coldest quarter (B). Gray shading represents 95% confidence intervals.

